# Identification of Circular RNAs associated with Ageing of the Dorsolateral Prefrontal Cortex across the Adult Lifespan

**DOI:** 10.1101/2024.12.10.627656

**Authors:** Fatemeh Amjadi-Moheb, Sumangali Gobhidharan, Adith Mohan, Perminder S. Sachdev, Anbupalam Thalamuthu, Karen A Mather

## Abstract

**Background:** Circular RNAs (circRNAs) are emerging as crucial regulators of biological processes and have been implicated in age-related diseases. Few studies have explored age-related circRNA expression in the human brain across the adult lifespan. This study aims to identify age-related differentially expressed circRNAs in human post-mortem dorsolateral prefrontal cortex (DLPFC) samples, a region critically involved in cognition that exhibits early signs of age-related changes.

**Methods:** Total RNA sequencing was conducted on a discovery cohort of 67 postmortem DLPFC samples, from individuals with no neurological disease diagnosis at the time of death (35-103 years). CircRNAs were identified using CIRCexplorer2, with 11,907 circRNAs available for analyses. Linear regression was used to analyse the relationships between circRNA expression and age at death. Replication of the results was performed in an independent neurologically healthy dataset from the CommonMind Consortium (n=321, age at death: 35-91 years). Co-expression network analysis was performed to identify modules of highly co-expressed circRNAs associated with age. Potential microRNA and RNA-binding protein target sites were predicted.

**Results:** In the discovery dataset, 37 circRNAs were age-associated (FDR <0.05). Seven out of the 37 were successfully replicated. The host genes of replicated age-associated circRNAs are implicated in synapse regulation. Co-expression analysis revealed two circRNA modules significantly correlated with age. We identified 484 microRNA and 99 RNA-binding protein target sites on the replicated circRNAs.

**Conclusion:** Seven age-associated circRNAs were identified as important candidates for involvement in post-transcriptional regulatory networks in the DLPFC. Future studies should aim to elucidate their functional roles in brain ageing.

## Introduction

Recent research has highlighted the pivotal role of circular RNAs (circRNAs) in various biological processes, particularly in the context of neurodegenerative disorders and age-related changes in the brain [1, 2]. CircRNAs are a class of RNA molecules, produced by the back splicing of exon and/or introns and characterised by their covalently closed-loop structure. They have been implicated in the regulation of gene expression through diverse mechanisms, including acting as microRNA (miRNA) and RNA-binding protein (RBP) sponges and influencing alternative splicing [3, 4].

CircRNAs are enriched in brain tissues and exhibit distinct expression patterns across different brain regions and developmental stages of different species [5–7]. The emerging importance of circRNAs in brain function is gaining significant support, particularly in the context of ageing and neurodegeneration [2, 8, 9].

Several studies show an age-related accumulation of circRNAs in the brain in *Drosophila*, *Caenorhabditis elegans*, mice, rats, pigs, monkeys, and humans [10–16]. However, to date, few studies have looked at the circRNA expression changes in the human brain across the lifespan [13, 17–19]. Liu et al. analysed a large dataset (n= 589) comprised of 312 psychiatric patients and 277 ‘healthy’ controls, revealing numerous age-associated circRNAs (n=175) in the dorsolateral prefrontal cortex (DLPFC), including well-known circRNAs like *CDR1as* [13]. In contrast, Gokool et al. [17] (n=144) and Cervera-Carles et al. [18] (n=69) observed no age-related trends in the cerebral cortex, frontal cortex, or cerebellum, while a study in the Netherlands noted circRNA accumulation with age in the substantia nigra (n=27) [19]. Methodological differences may have contributed to the inconsistencies of the observed results. It is worth noting that the majority of these studies looked at age as a covariate in the analysis rather than focusing directly on circRNA expression changes with age.

Significant methodological variations and limited sample sizes in these studies emphasise the necessity for additional research to clarify circRNA dynamics in healthy brain ageing. The DLPFC is a key region in networks subserving executive functions such as planning, decision-making, and problem-solving and is vulnerable to ageing [20, 21]. It exhibits a decline in volume and sensitivity to age-related morphological and functional changes such as decreased activation patterns during cognitive tasks, compensatory recruitment of posterior brain regions, and declines in performance on tasks involving executive functions and memory [22, 23]. This study aims to provide a comprehensive analysis of circRNA expression profiles in the ageing DLPFC across adulthood.

## Materials and methods

### Study Cohorts

#### Discovery cohort (Brain Ageing Study)

This study used a discovery cohort from the Brain Ageing Study, comprising 67 post-mortem brain samples derived from the DLPFC (Brodmann areas 46, 9, or 10). These samples were selected from individuals devoid of neurological pathologies. Spanning an age range of 35.6 to 103 years, the samples were acquired from two repositories: the Human Brain Collection Core (HBCC) within the National Institute of Mental Health’s (NIMH) Division of Intramural Research Programs (n=41, mean age 56 years), and the Sydney Brain Bank (SBB; n=26, mean age 91 years). The Brain Ageing study received approval from the local Human Research Ethics Committee of the University of New South Wales.

#### Replication Dataset (a subset of the CommonMind Consortium [CMC] dataset)

The replication cohort data from control samples free of neurological disease [24] were retrieved from an independent RNA-seq dataset previously collected by the CommonMind Consortium (CMC) (n=321). This cohort spans an age range of 35 to 90.5 years, matching the age range of the discovery dataset. The replication dataset encompasses samples from four different brain banks: the Mount Sinai NIH Brain and Tissue Repository (n=139, age at death: 36-90.5 years), the University of Pennsylvania Alzheimer’s Disease Core Center (n=35, age at death: 36-90.5 years), the University of Pittsburgh NeuroBioBank and Brain and Tissue Repositories (n=71, age at death: 36-82 years), and the NIMH HBCC (n=76, age at death: 35-73 years).

### RNA Sequencing and CircRNA Prediction

#### Discovery Study

Total RNA was extracted from approximately 30 mg of brain tissue using the RNeasy mini kit (Qiagen, Toronto, ON, Canada) according to the manufacturer’s protocol, followed by treatment with 1 μl DNase I (Qiagen). After quality control assessments, rRNA-depleted cDNA library preparation (Illumina TruSeq Stranded Ribozero, Illumina Inc, San Diego, CA, USA) was conducted at the UNSW Ramaciotti Centre for Genomics, and sequencing was performed on an Illumina NovaSeq S4 sequencer to obtain 100-bp paired-end reads with a minimum of 50 million reads per sample. The raw data underwent preprocessing procedures utilising Trim Galore v.0.6.10 [25] to remove adapter sequences and sequences of low quality, followed by an assessment of data quality using FastQC [26]. Subsequently, all FASTQ files underwent alignment to the human reference genome (hg38, Ensembl v97) employing STAR 2.7.6a [27]. Further analysis involved the identification of circRNAs utilising CIRCexplorer2 [28], employing default parameters suitable for paired-end RNA-seq data. Another tool, CIRI2 [29], was used for comparison with CIRCexplorer2 (see supplementary document). To ensure data integrity, circRNAs were determined to be expressed only if they exhibited back-spliced junctions in over 50% of the samples, with a minimum of two junction reads per circRNA. Subsequently, circRNA expression levels were normalised to count per million reads (CPM), and log2 transformed to mitigate the impact of library depth variations.

### Replication Data Set

Replication was undertaken using the CMC dataset, which had RNA sequencing previously performed, and the BAM format files were obtained from the Synapse platform at http://CommonMind.org, and subsequently merged and converted to FASTQ format utilising SAMtools (v1.19.1) [30]. Notably, samples from the NIMH HBCC were generated utilising a stranded library protocol, while other samples within the CMC dataset were generated using a non-stranded library protocol. Except for trimming, the processing steps applied to the CMC dataset samples remained consistent, accounting for the RNA-seq library protocol variations (supplementary methods).

### Statistical Analyses

All statistical analyses and data visualisation were performed in R v4.3.1 (R Foundation, Vienna, Austria). A linear regression model was applied to analyse the relationship between age and expression of circRNAs on log2 transformed counts per million reads (CPM), adjusting for potential confounders (sex, postmortem intervals, estimated proportion of neuronal cell counts (see supplementary document), RNA integrity (RIN), pH as well as library protocol type for samples of CMC dataset). For all tests, Benjamini-Hochberg (BH) correction for multiple testing was performed and a false discovery rate (FDR) value < 0.05 was considered statistically significant.

### Functional and Pathway Enrichment Analyses

We performed Gene Ontology (GO) functional enrichment analysis of the parental genes of significant circular RNAs using the enrichR R package [31], using the SynGO database [32], which is a specialised resource for brain-related gene sets. An adjusted p-value < 0.05 significance threshold was applied to identify enriched GO terms.

### CircRNA Co-expression Network Analyses

The co-expression network analysis was performed on the discovery dataset using a signed network approach implemented in the WGCNA (v1.72.1) R package [33]. Detailed methods are described in the supplementary methods.

### Prediction of binding sites for miRNAs and RNA-binding Proteins

To elucidate the regulatory effects of the age-associated circRNAs, we predicted putative binding sites for miRNAs and RNA-binding proteins on the seven replicated and age-related circRNAs. The sequence of age-related circRNAs were obtained from circAtlas database [34]. For miRNA binding site prediction on circRNA sequences, we employed two alternative algorithms, miRanda 3.3a [35] and PITA [36], utilising specific parameters for miRanda: score_miRNA=135 and default settings for PITA. Only miRNA binding sites predicted concurrently by false discovery rate with both algorithms were retained for further analysis. miRNA-mRNA interactions were predicted using three established miRNA databases (TargetScan [37], miRWalk [38], and miRTarBase [39]), retaining only those interactions predicted by all databases to ensure robustness. Subsequently, based on the predicted binding sites, we constructed and visualised a circRNA–miRNA–mRNA network for the top results, using Cytoscape software (v3.10.1) [40].

For RBP binding site prediction, we utilised the beRBP [41] algorithm with the beRBP general model to predict RBP binding sites within the significant circRNAs, filtering for those with a beRBP score greater than or equal to 0.43, which represents a cutoff point chosen to prioritise high-confidence binding site predictions while maintaining a reasonable number of candidate sites for downstream analysis.

## Results

Detailed demographic summaries of the Brain Ageing and CMC datasets are provided in Table 1, indicating that the discovery dataset consists of an older population (mean age at death = 71.40 ± 20.70) compared to the replication cohort (mean age at death = 62.00 ± 17.01). In the original CMC dataset, individuals aged 90 and above were coded as 90+, but for the purposes of our analysis, they were reassigned as 90.5.

**Table 1.** Demographic characteristics of participants.

| **Characteristics** | **Brain Ageing Study**<br>**(Discovery Dataset)** | **CMC Study**<br>**(Replication Dataset)** |
| --- | --- | --- |
| **Participants, N** | 67 | 321 |
| **Age at Death Range, Years** | 35.6-103 | 35- 90.5* |
| **Mean Age, Years (SD)** | 71.40 (20.70) | 62 (17.01) |
| **Sex: Female, N (%)** | 30 (45) | 132 (41) |
| **Mean PMI, Hours (SD)** | 25.50 (15.64) | 17.30 (11.87) |
| **Neuron Percentage (%)** | 44 | 44 |
| **Mean RIN Score (SD)** | 5.60 (1.52) | 7.90 (0.89) |
| **Mean pH (SD)** | 6.39 (0.29) | 6.57 (0.26) |

\* In the CMC dataset, individuals aged greater than 90 were assigned an age of 90.5. SD: standard deviation; N: number; PMI: postmortem interval; RIN: RNA integrity.

Despite the lower RNA Integrity Number (RIN) in the discovery RNA-seq dataset (mean RIN = 5.60 ± 1.52) compared to the replication cohort (mean RIN = 7.90 ± 0.89), the RNA-seq quality control (QC) metrics, including high alignment rates and consistent sequencing depth, indicated robust results.

### Identification of circRNAs

A total of 11,907 circRNAs within the discovery and replication datasets using CIRCexplorer2 were detected. The exons in the identified circRNAs ranged from 1 to ≥10, with approximately half of the identified circRNAs containing 2–4 exons (51.83%; Figure 1.a). The median length of detected circRNAs was 376 base pairs (bp) (Figure 1.b). The distribution of circRNAs (n=11,907) across parental genes (n=3,870), showed that the majority of genes (95.56%) yielded one to ten distinct circRNAs. However, some parental genes (e.g. *ATRNL1*, *PTK2*, *ADGRB3*), generated more than 40 unique circRNAs (circRNA isoforms; Figure 1.c). In the replication cohort (Figure S1), a similar pattern was observed.

**Figure 1.**
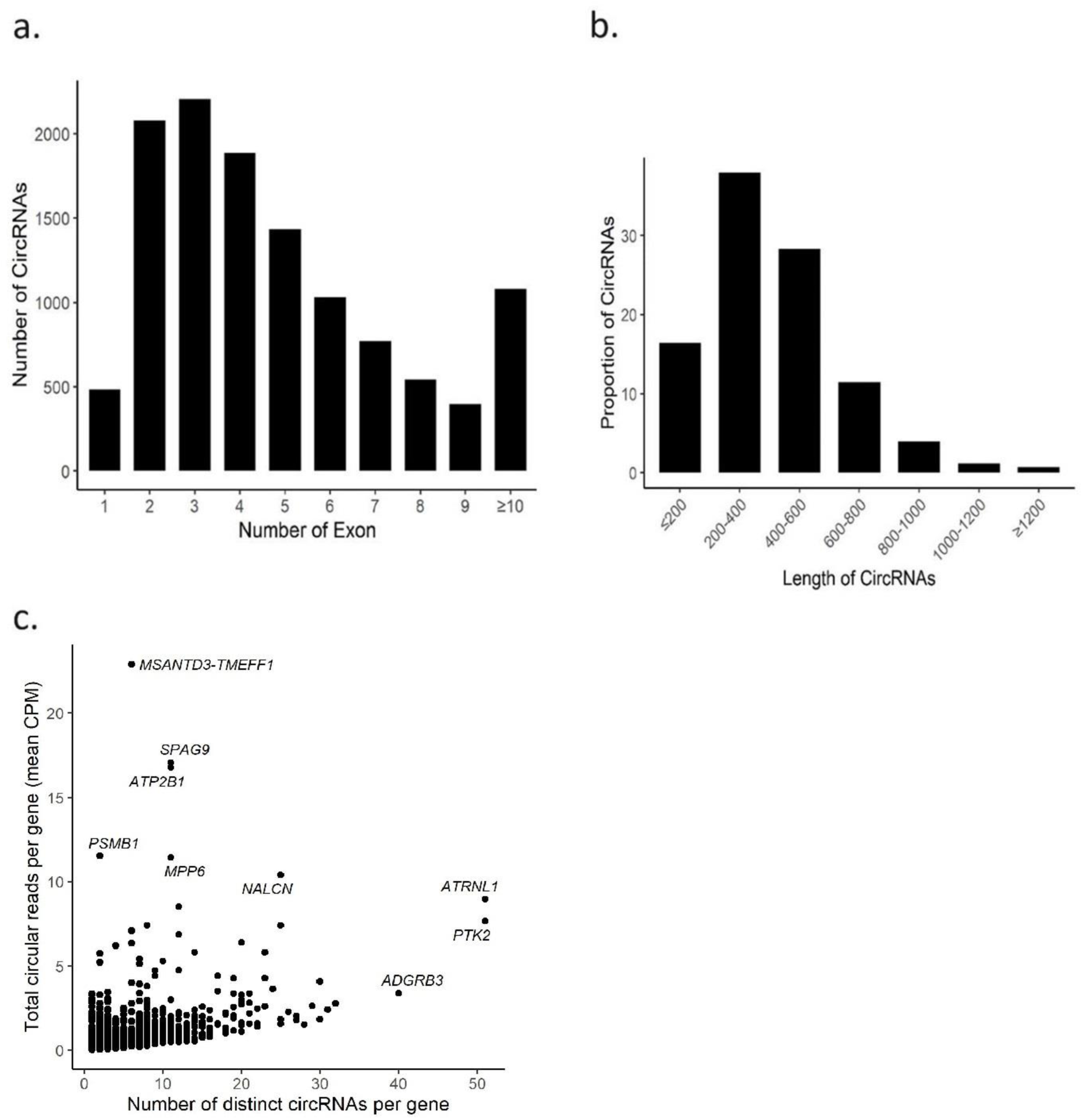
CircRNA profile in human brain ageing across the lifespan. **a.** Distribution of circRNA exons. **b.** Distribution of circRNA lengths. **c.** Comparison of the mean CPM of circular reads per gene with the number of distinct circRNAs per gene in the discovery dataset.

### Differential Expression of circRNAs Across Adult Lifespan

Using a linear regression model, in the discovery dataset, 37 circRNAs out of 11,907 exhibited significant differential expression patterns associated with age (FDR < 0.05; Table S1). The majority were upregulated with age, with only 11 downregulated. The top downregulated circRNA was *HOMER1_0006*, derived from the Homer Scaffold Protein 1 (*HOMER1*) gene, and the top upregulated circRNA, *MIR31HG_0001*, was derived from the *MIR31HG* gene, a long non-coding RNA that also encodes miR-31. All 37 circRNAs were available in the CMC dataset for replication. Seven were differentially expressed in the same direction with age and were of similar effect size in the replication CMC dataset (Table 2; Figure 2). These replicated and age-associated circRNAs were derived from the following parental genes, *HOMER1*, neuropilin and tolloid-like 2 (*NETO2*), special AT-rich sequence-binding protein 1 (*SATB1*), RAS guanyl releasing protein 1 (*RASGRP1*), thyroid hormone receptor interacting protein 12 (*TRIP12*), and LIM domain 7 (*LMO7*). Notably, two circRNAs were derived from the *HOMER1* gene. The circRNA *TRIP12_0034* was up-regulated with age, while *HOMER1_0007, LMO7_0001, HOMER1_0006, NETO2_0001, SATB1_0015*, and *RASGRP1_0003* were down-regulated with age.

**Table 2.** Replicated age-associated circRNAs in the DLPFC.

| CircRNA | Chr | Start\| End (Strand) | Discovery dataset |  |  | Replication dataset |  |  |
| --- | --- | --- | --- | --- | --- | --- | --- | --- |
|  |  |  | Beta | SE | FDR | Beta | SE | p-value |
| *HOMER1_0006* | 5 | 79439010\|79457018(-) | -0.015 | 0.002 | 1.01E-05 | -0.018 | 0.002 | 5.95E-16 |
| *NETO2_0001* | 16 | 47109483\|47132025(-) | -0.013 | 0.003 | 0.01 | -0.010 | 0.002 | 1.14E-05 |
| *SATB1_0015* | 3 | 18378170\|18420991(-) | -0.014 | 0.003 | 0.01 | -0.007 | 0.002 | 0.004 |
| *RASGRP1_0003* | 15 | 38502312\|38505920(-) | -0.017 | 0.003 | 0.015 | -0.018 | 0.003 | 7.35E-08 |
| *TRIP12_0034* | 2 | 229774097\|229788940(-) | 0.017 | 0.004 | 0.021 | 0.010 | 0.003 | 1.1E-03 |
| *HOMER1_0007* | 5 | 79439010\|79451121(-) | -0.017 | 0.004 | 0.028 | -0.023 | 0.003 | 6.48E-13 |
| *LMO7_0001* | 13 | 75713182\|75761038(+) | -0.016 | 0.004 | 0.042 | -0.009 | 0.003 | 0.002 |

A linear regression model was used to assess the relationship between circRNA expression and age, adjusting for relevant covariates. Chr: Chromosome; Beta: Estimated coefficient for age in the linear regression. SE: Standard error associated with the Beta coefficient; FDR: False discovery rate.

**Figure 2.**
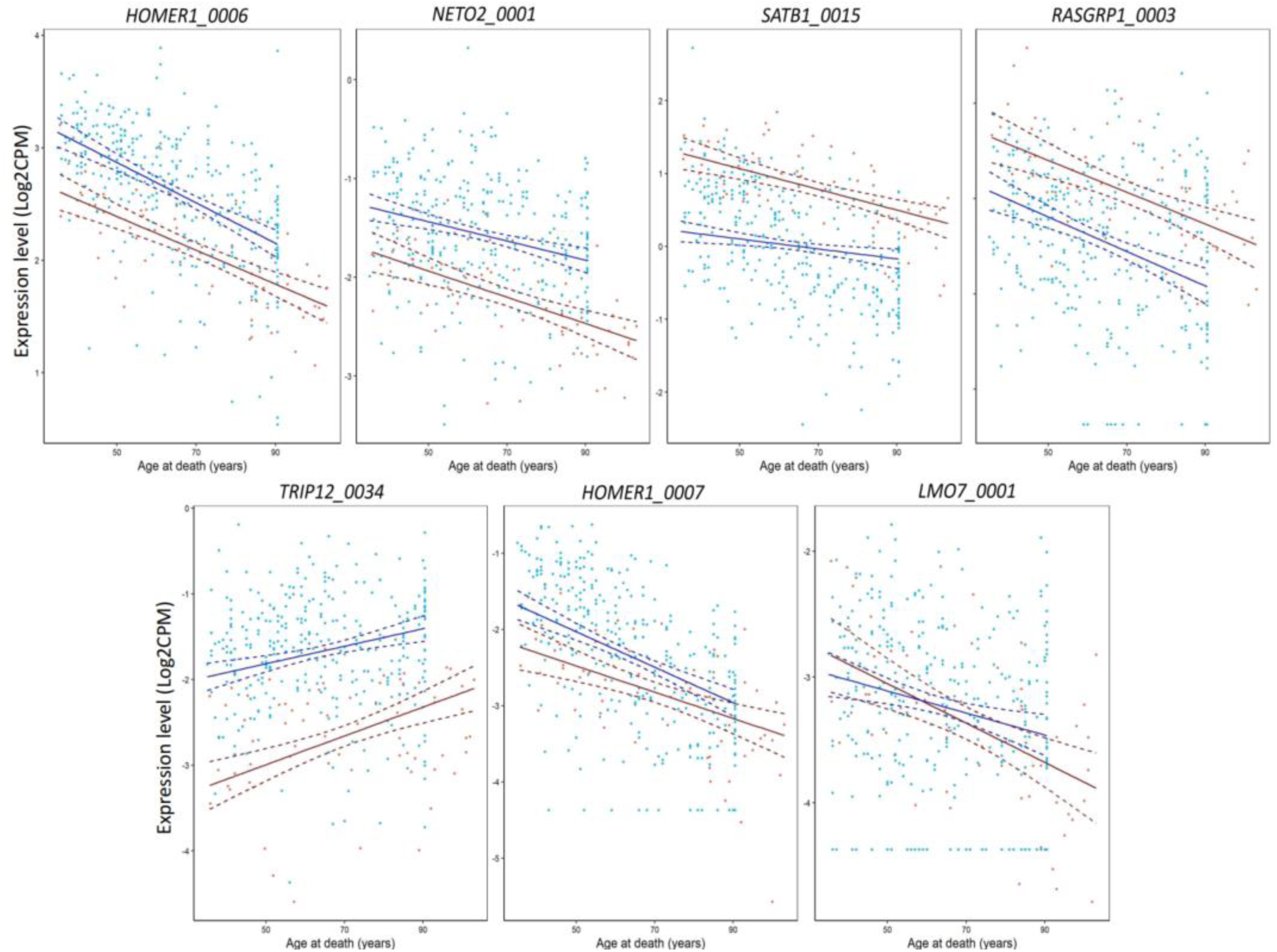
The expression of replicated circRNAs with age at death in both datasets. Red: discovery dataset; Blue: replication dataset.

### CircRNA Co-expression Network Analyses

The co-expression network analysis identified 10 distinct modules of highly co-expressed circRNAs in the discovery dataset. One module (grey module) contained circRNAs that could not be assigned to any specific module (see supplementary document). Pearson correlation coefficients (r) were used to evaluate the association between each module and age at death. Among the 10 modules, module M4 exhibited a significant positive correlation with age (n=473; r = 0.41, p = 5e-04), while module M5 showed a significant negative correlation (n=413; r = −0.32, p = 0.007, adjusted p-value = 0.005 for multiple testing; Figure 3). Notably, both modules contained circRNAs identified as age-related in the discovery dataset, with module M5 also including replicated age-related circRNAs: *HOMER1_0006, SATB1_0015, RASGRP1_0003,* and *LMO7_0001*. Functional enrichment analysis of the parental genes of circRNAs in modules M4 and M5 revealed enrichment for genes involved in synaptic vesicle priming, synaptic vesicle endocytosis, and other critical processes related to synaptic activity (Figure S2).

**Figure 3.**
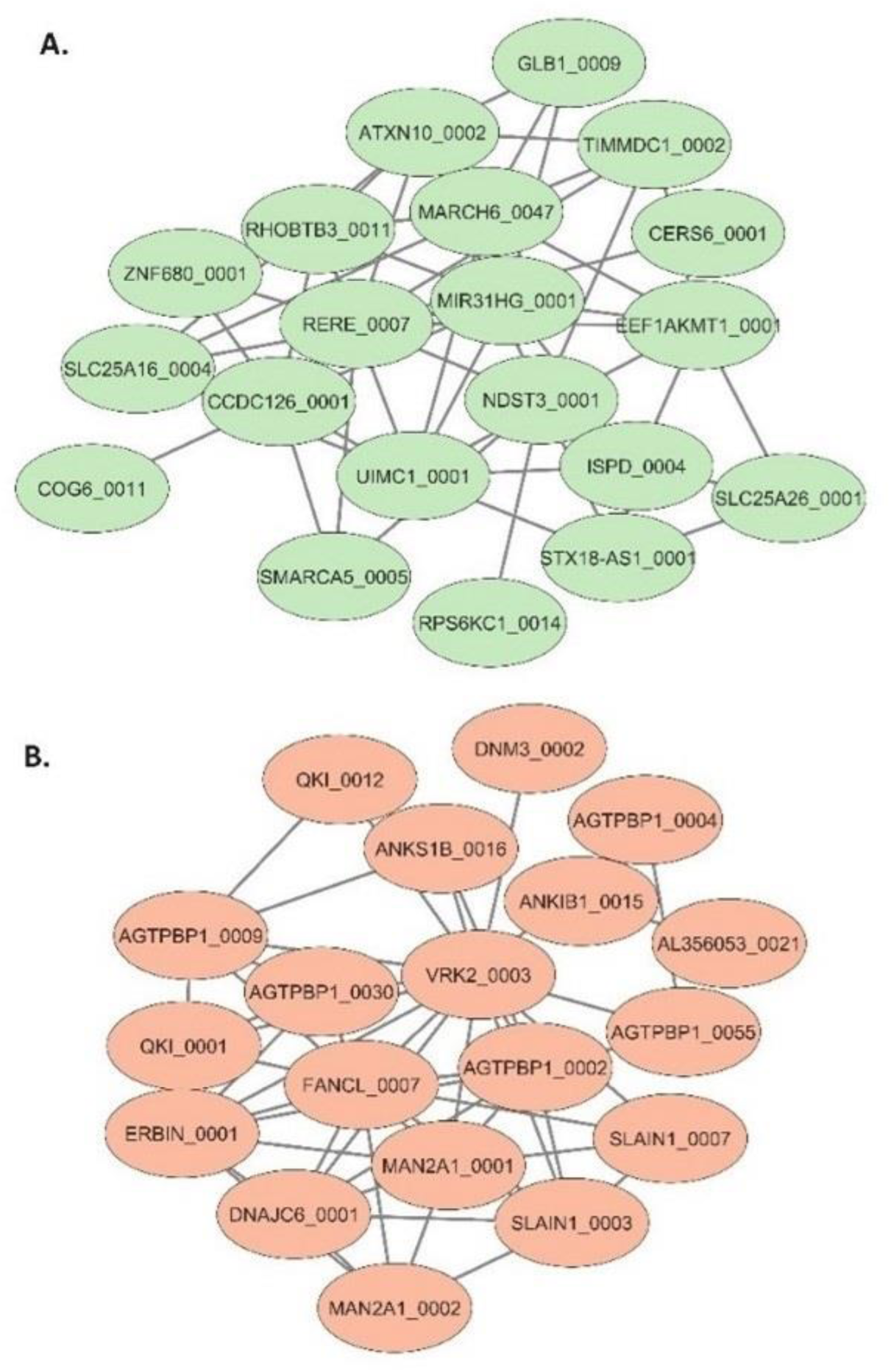
**A.** M4 Module and **B.** M5 Module: The top 20 circRNAs with the highest connectivity scores within this module are visualised using Cytoscape. Each node represents a circRNA, and edges indicate co-expression relationships.

### Functional Insights of CircRNAs Throughout the Lifespan

Functional enrichment analysis of the host genes of the 37 age-associated circRNAs in the discovery dataset revealed significant Gene Ontology (GO) terms primarily involved in synapse regulation. The top enriched biological processes (BP) included the regulation of postsynaptic neurotransmitter receptor activity (GO:0098962; FDR = 0.017) and the regulation of neurotransmitter receptor localisation to the postsynaptic specialisation membrane (GO:0098696; FDR = 0.019; Table S2).

### Identification of miRNA and RBP Binding Sites in Age-Related circRNAs and Construction of a Regulatory Network

To identify miRNAs potentially regulated by the replicated age-related circRNAs, we bioinformatically predicted miRNA binding sites in circRNA sequences (Table S3). As two age-related circRNA isoforms of *HOMER1* were identified, for this analysis, the longer isoform, *HOMER1_0006*, was included because it completely encompasses the sequence of the shorter isoform (*HOMER1_0007*). A total of 484 unique miRNAs were predicted to bind to at least one of the age-related circRNAs. *HOMER1_0006* contains binding sites for 110 miRNAs, with 80 out of the 484 predicted miRNAs binding exclusively to *HOMER1_0006*. In contrast*, TRIP12_0034* has the fewest miRNA binding sites among the age-related circRNAs, with binding sites for 63 miRNAs, of which 46 miRNAs bind uniquely to *TRIP12_0034*.

Our study showed that the 484 miRNAs can potentially bind up to three age-related circRNAs each. Figure 4 shows the six miRNAs that had binding sites on three circRNAs. The mRNA targets of these six miRNAs were identified bioinformatically, and a circRNA–miRNA–mRNA regulatory network was constructed based on these interactions (Figure 4).

**Figure 4.**
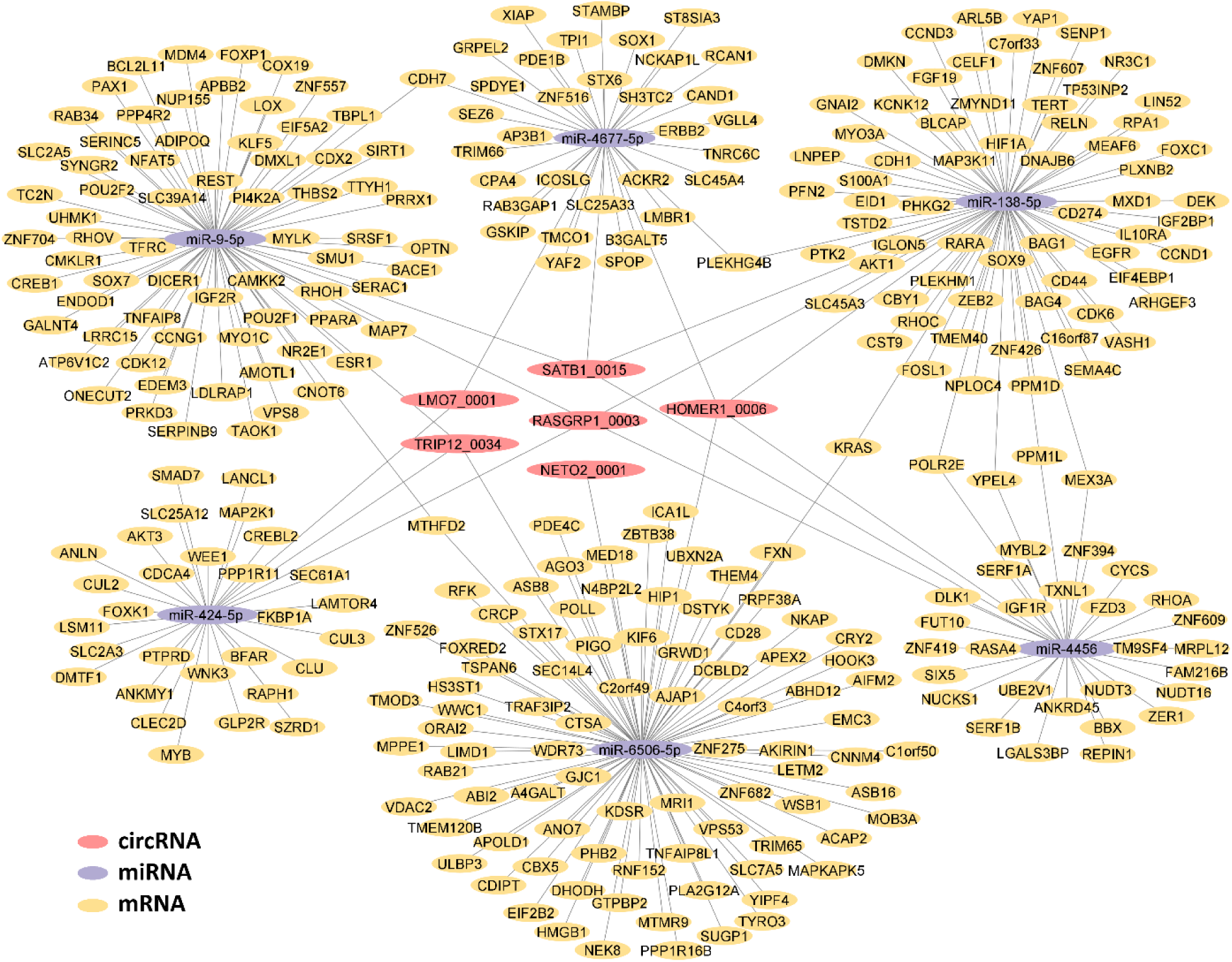
The circRNA–miRNA–mRNA network of the age-associated replicated circRNAs and their target miRNAs and mRNAs. This network focuses on the six miRNAs (out of the 484 predicted miRNAs) identified as having binding sites on three different age-related circRNAs. These six miRNAs and their bioinformatically predicted mRNA targets are depicted, while miRNAs that bind to two or fewer circRNAs are not shown.

Additionally, we identified potential RBP binding motifs within the age-related circRNAs (Table S4). A total of 99 RBP binding sites were predicted across the age-related circRNAs, with *HOMER1_0006* containing 80 of these sites, while *LMO7_0001* had the fewest, with only 9 sites. Notably, splicing factors and neural-specific RBPs, such as Serine and Arginine-Rich Splicing Factor 1 (SRSF1) and Eukaryotic Translation Initiation Factor 4B (EIF4B), were among the top results, with multiple predicted binding sites on the circRNAs, except for *LMO7_0001*.

## Discussion

In this study, we aimed to identify age-related differentially expressed circular RNAs in the DLPFC region collected from deceased individuals aged 35-103 years. In the discovery analysis, we observed 37 circRNAs associated with age of death. Seven of these were subsequently replicated in an independent cohort. The majority of the replicated age-associated circRNAs demonstrated downregulation with advancing age, namely *HOMER1_0006, NETO2_0001, SATB1_0015, RASGRP1_0003, HOMER1_0007,* and *LMO7_0001*, while one circRNA, *TRIP12_0034*, was upregulated.

Previous studies indicate that the replicated circRNAs derived from *HOMER1* and *SATB1* play roles in synaptic plasticity, signal transduction, and neuronal homeostasis, and have been implicated in various neurological and psychiatric disorders. For instance, circRNAs from the *HOMER1* gene, are associated with cognitive function and neurodegeneration, and their downregulation is linked to Alzheimer’s disease, Parkinson’s disease, schizophrenia, and bipolar disorder [18, 42–47]. *SATB1_0015* has been shown to influence dendritic spine morphology, with its decreased expression resulting in dendritic abnormalities in mesial temporal lobe epilepsy [48]. Their parental genes, *HOMER1* and *SATB1*, contribute to critical brain functions, such as chromatin organization, neuronal migration, and synaptic connectivity, and their dysregulation has been implicated in conditions like Parkinson’s disease, epilepsy, and schizophrenia [49–56].

Although there is no current evidence regarding the functional roles of the other replicated circRNAs (*LMO7_0001, NETO2_0001, RASGRP1_0003,* and *TRIP12_0034*), their parental genes play crucial roles in neurological functions and disorders. *LMO7* is associated with psychotic disorders and temporal lobe epilepsy [57–60], *NETO2* enhances synaptic plasticity and neurotransmission through kainate receptor interactions [61–64], *RASGRP1* is implicated in bipolar disorder and memory formation [65–69], and *TRIP12* influences neuroinflammation and Parkinson’s pathology via protein regulation [70, 71]. Collectively, these results highlight the significant roles of parental genes in neuropsychiatric disorders and brain ageing.

Co-expression network analysis identified two modules of co-expressed circRNAs that exhibited significant correlations with age at death in the discovery dataset, one positively and one negatively. Most of the age-associated circRNAs in the discovery dataset were found within these modules. Notably, the module with a negative correlation to age contained several replicated age-related downregulated circRNAs, including *HOMER1_0006, SATB1_0015, RASGRP1_0003*, and *LMO7_0001*. Functional enrichment analysis of the parental genes of circRNAs in these two modules revealed key roles in synaptic vesicle priming, synaptic vesicle endocytosis, and synaptic activity, suggesting their involvement in preserving synaptic function during ageing.

Through bioinformatic analysis, we identified potential binding sites for miRNAs and RBPs on circRNAs, elucidating candidate regulatory networks involved in age-related processes. Among the identified miRNAs, miR-424 has been shown to be upregulated in Alzheimer’s disease and plays a role in Parkinson’s disease, where it influences energy metabolism by targeting genes related to fatty acid synthesis [72–74]. Similarly, miR-138-5p is crucial in regulating neuronal processes like dendritic spine morphology [75, 76] and synaptic plasticity [77, 78], with its dysregulation linked to Alzheimer’s disease and amyloid-beta (Aβ) production, underscoring its impact on synaptic health and cognitive decline during ageing [76, 79, 80]. Additionally, the Serine/Arginine-rich Splicing Factor 1 (SRSF1) and Eukaryotic initiation factor 4B (EIF4B), identified as top RNA binding proteins (RBPs) bound to circRNAs, play crucial roles in post-transcriptional regulation and neuronal development [81–84]. The results underscore the potential significance of the identified age-associated circRNAs in brain function and health.

Previously, Liu et al. [11], used a sub cohort of the CMC dataset to look at the circRNA expression patterns in the DLPFC and found 175 age-related circRNAs. The Liu et al. subset comprised samples from both psychiatric disorder patients (schizophrenia and affective/mood disorders) and healthy controls across all age ranges, from infants to the oldest old. In comparison to their study, our investigation takes a different approach and focus, specifically targeting neurologically healthy control samples within a narrower age range to delve into age-associated circRNA expression changes in adults (age at death: 35-90 years). Interestingly, three of the seven age-associated circRNAs in the current study, *HOMER1_0006, NETO2_0001,* and *RASGRP1_0003*, showed consistent downregulation trends in both the Liu et al. study and ours, suggesting potential robustness across different datasets and study designs. Notably, our study extends Liu et al.’s findings by focusing on age-related changes in circRNA expression within a healthy ageing context, offering complementary insights into the potential role of circRNAs in age-related processes in the DLPFC.

Two studies have not found age-related differences in circRNA expression [17, 18], and the discrepancies between our results and their findings may be attributed to methodological differences. First, variations in sequencing platforms and bioinformatics pipelines might have influenced circRNA detection sensitivity. For example, Gokool et al. [17] used unstranded ribosomal RNA-depleted RNA-seq, which may lack the specificity required for identifying subtle age-related expression changes. Cervera-Carles et al. [18] employed RT-qPCR, which, while precise for targeted validation, is less comprehensive than RNA-seq for profiling the full circRNA transcriptome. Second, smaller sample sizes or differences in demographic characteristics, such as varying age distributions, might have reduced the power of these studies to identify age-related patterns. Third, these studies primarily focused on neurodegenerative conditions, potentially masking age-related trends in healthy individuals [17, 18]. Finally, another possible reason for the discrepancy is the specific brain regions examined in these studies. Differences in circRNA expression patterns across brain regions may contribute to the observed inconsistencies. Cervera-Carles et al. [18] focused on the frontal cortex, while Gokool et al. [17] studied the frontal cortex, temporal cortex, and cerebellar vermis. In contrast, the present study was conducted exclusively in the dorsolateral prefrontal cortex (DLPFC), a region with distinct structural and functional properties.

This study represents an exploration of circRNA expression in the DLPFC across a wide age range in adults free of neurological disease diagnosis at the time of death. Unlike previous research, which often focused on neurodegenerative disease with limited age ranges, this investigation offers an important foundation for understanding how circRNAs change with ageing in neurologically healthy human brain samples. However, the study has some limitations. One limitation is the reliance on total RNA-seq data and bioinformatic approaches for circRNA identification. While these methods offer valuable insights, experimental validation techniques, such as RT-qPCR, were not used to confirm the expression patterns of the identified circRNAs. However, we did use an independent cohort for replication of the discovery results.

Future studies could utilise multiple ‘omic assays, such as performing standard RNA and small RNA sequencing on the same sample. This approach would enable more comprehensive, data-driven analyses rather than solely relying on bioinformatic predictions. A comparative analysis of age-related circRNA trajectories in other brain regions, particularly those that exhibit early age-related changes (e.g., the posterior cingulate cortex), would be of significant interest. This will be the focus of our future investigations.

## Conclusion

In summary, our study provides a comprehensive analysis of circRNA expression profiles in the ageing DLPFC, revealing age-associated changes in circRNA expression across the adult lifespan. The identification of age-associated circRNAs in the DLPFC offers valuable insights into potential regulatory networks underlying brain ageing and provides targets for further work. Future research could further investigate the specific circRNAs identified here, exploring their roles, interactions with other molecules, and their potential use as diagnostic biomarkers or therapeutic targets in brain ageing.

## Supporting information

Supplementary analyses

## Acknowledgements

The authors express their deepest gratitude to the brain sample donors, their families, and the research teams of the Human Brain Collection Core (HBCC), Intramural Research Program, National Institute of Mental Health (NIMH), the Sydney Brain Bank (SBB), and the CommonMind Consortium (CMC). Tissue samples were obtained from the HBCC, NIMH, and the Sydney Brain Bank (SBB), which is supported by Neuroscience Research Australia (NeuRA). The authors gratefully acknowledge the CMC for providing access to the human dorsolateral prefrontal cortex (DLPFC) RNA-seq data. Data were generated as part of the CMC, supported by funding from Takeda Pharmaceuticals Company Limited, F. Hoffman-La Roche Ltd, and NIH grants R01MH085542, R01MH093725, P50MH066392, P50MH080405, R01MH097276, RO1-MH-075916, P50M096891, P50MH084053S1, R37MH057881, AG02219, AG05138, MH06692, R01MH110921, R01MH109677, R01MH109897, U01MH103392, and contract HHSN271201300031C through IRP NIMH. Brain tissue for the study was obtained from the following brain bank collections: the Mount Sinai NIH Brain and Tissue Repository, the University of Pennsylvania Alzheimer’s Disease Core Center, the University of Pittsburgh NeuroBioBank and Brain and Tissue Repositories, and the NIMH Human Brain Collection Core. We acknowledge the leadership of the CMC, including Panos Roussos, Joseph Buxbaum, Andrew Chess, Schahram Akbarian, and Vahram Haroutunian (Icahn School of Medicine at Mount Sinai); Bernie Devlin and David Lewis (University of Pittsburgh); Raquel Gur and Chang-Gyu Hahn (University of Pennsylvania); Enrico Domenici (University of Trento); Mette A. Peters and Solveig Sieberts (Sage Bionetworks); and Thomas Lehner, Stefano Marenco, and Barbara K. Lipska (NIMH). The analyses and conclusions presented in this study are those of the authors and do not necessarily reflect the views of the CMC, its funders, or any of the contributing institutions.

## Author Contribution

KM, AT, FA, & SG: designed the study and interpreted the results. KM, SG, & AM: provided samples and wet lab experiments. FA: research, collection, assembly of data, and manuscript writing and prepared figures. AT & FA: formed the database of samples, analysed data, & performed statistical analysis. PS & KM: provided financial support. KM: supervised the study. All authors gave input to the drafted manuscript and approved the final version for submission

## Data Availability

Due to ethics constraints only summary results can be shared. Additional details can be obtained from the authors upon reasonable request.

## Funding

This study was supported by Rebecca L. Cooper Medical Research Foundation Project Grant (KM).

## Declarations

### Ethics Approval

This study received approval from the local Human Ethics Committee of the University of New South Wales (UNSW). The research utilized human brain tissue and data provided by the Human Brain Collection Core (HBCC) of the National Institute of Mental Health (NIMH) Intramural Research Program, the Sydney Brain Bank (SBB) at Neuroscience Research Australia (NeuRA), and the CommonMind Consortium (CMC). Ethics approval for the use of human tissue was obtained from the respective institutions. The HBCC complies with the Federal Policy for the Protection of Human Subjects (Common Rule), and all research involving these tissues was approved by an Institutional Review Board (IRB). The Sydney Brain Bank provided tissue samples in accordance with ethics approvals granted by relevant ethics committees, following the Sydney Brain Bank Guidelines for Researchers. The RNA-seq data from the CMC were used under the ethical oversight of the respective IRBs of participating institutions.

### Consent to Participate

All human brain tissue used in this study was collected with informed consent from the donors or their legal representatives. Donors provided consent for the use of their tissue and data in scientific research to advance understanding of brain function and disorders. Strict measures were implemented to ensure the confidentiality and privacy of all donors throughout the study.

## Notes

### Competing Interest Statement

The authors have declared no competing interest.

