## Supplementary analyses for "Identification of Circular RNAs associated with Ageing of the Dorsolateral Prefrontal Cortex across the Adult Lifespan"

**Methods:**

**Identification of circRNAs from RNA-seq Data**

After trimming, the FASTQ reads of discovery and replication RNA-seq datasets were aligned to the human reference genome (hg38, Ensembl v97) using STAR 2.7.6a [1], with optimised parameters for splice junction detection: --outSJfilterOverhangMin 15 15 15 15, --alignSJoverhangMin 15, --alignSJDBoverhangMin 15, --outFilterMultimapNmax 20, --outFilterScoreMin 1, --outFilterMatchNmin 1, --outFilterMismatchNmax 2, --chimSegmentMin 15, --chimScoreMin 15, --chimScoreSeparation 10, and --chimJunctionOverhangMin 15. For samples with the non-stranded library protocol (CMC data obtained from the Mount Sinai NIH Brain and Tissue Repository, the University of Pennsylvania Alzheimer’s Disease Core Center, the University of Pittsburgh NeuroBioBank, and Brain and Tissue Repositories), we also included the --outSAMstrandField intronMotif option to ensure accurate alignment.

Following alignment, CIRCexplorer2 was run with its default parameters for paired-end data, applying specific library-type modifications for stranded and non-stranded library preparation protocols. In addition, CIRI2 [2], was utilised, using the BWA aligner for initial read alignment with default parameters, adjusted according to the library preparation types.

**Cell Type Composition Analysis**

We estimated the proportions of individual cell types in our brain tissue samples using DeconRNAseq. For reference transcriptomes, we used gene expression data from purified cell populations (neurons, astrocytes, oligodendrocytes, microglia, and endothelial cells) provided by Zhang et al. [5].

**CircRNA Co-expression Network Analyses**

For the co-expression network analysis, the normalised circRNA expression values underwent covariate correction, accounting for sex, postmortem intervals, proportion of neuronal cell counts, RNA integrity (RIN), and pH, using a linear model. The residual values were then used for automatic network construction employing the `blockwiseModules` function with the following parameters: power = 11, networkType = "signed", corFnc = "bicor", minModuleSize = 30, and mergeCutHeight = 0.25.

**Results**

**Comparative Analysis of circRNA Detection Using CIRCexplorer2 and CIRI2**

In our comparative analysis using two circRNA identification tools, CIRCexplorer2 identified 11,907 circRNAs, while CIRI2 detected 14,429 circRNAs across the discovery cohort samples. Among the circRNAs detected, 3,790 were unique to CIRI2, while 1,268 circRNAs were uniquely identified by CIRCexplorer2. Notably, both tools identified 10,639 circRNAs in common, representing 91% of the total circRNAs detected, indicating a high degree of concordance between them (Figure S3. A). Furthermore, the correlation between circRNA expression levels identified by CIRCexplorer2 and CIRI2 within the same samples ranged from 0.91 to 0.97 (Figure S3. B and C), underscoring the robustness of circRNA quantification across these two tools. We chose CIRCexplorer2 to use in our analyses as the proportion of filtered circRNAs meeting our threshold criteria was notably higher (CIRCexplorer2 69% versus 35% for CIRI2 35%).

**Module Assignment of circRNAs in Co-Expression Network Analysis**

For the co-expression network analysis, the kME values for each circRNA are provided in Table S6. The module eigengene is the first principal component of the expression data for the circRNAs within that module and serves as a representative expression pattern for all the members of that module. The kME values represent the correlation between each circRNA and the module eigengene. CircRNAs were grouped into modules according to their correlation with the module eigengene (kME > 0.1) and the statistical significance of this correlation, determined by a BH-corrected p-value (FDR < 0.05) [4].

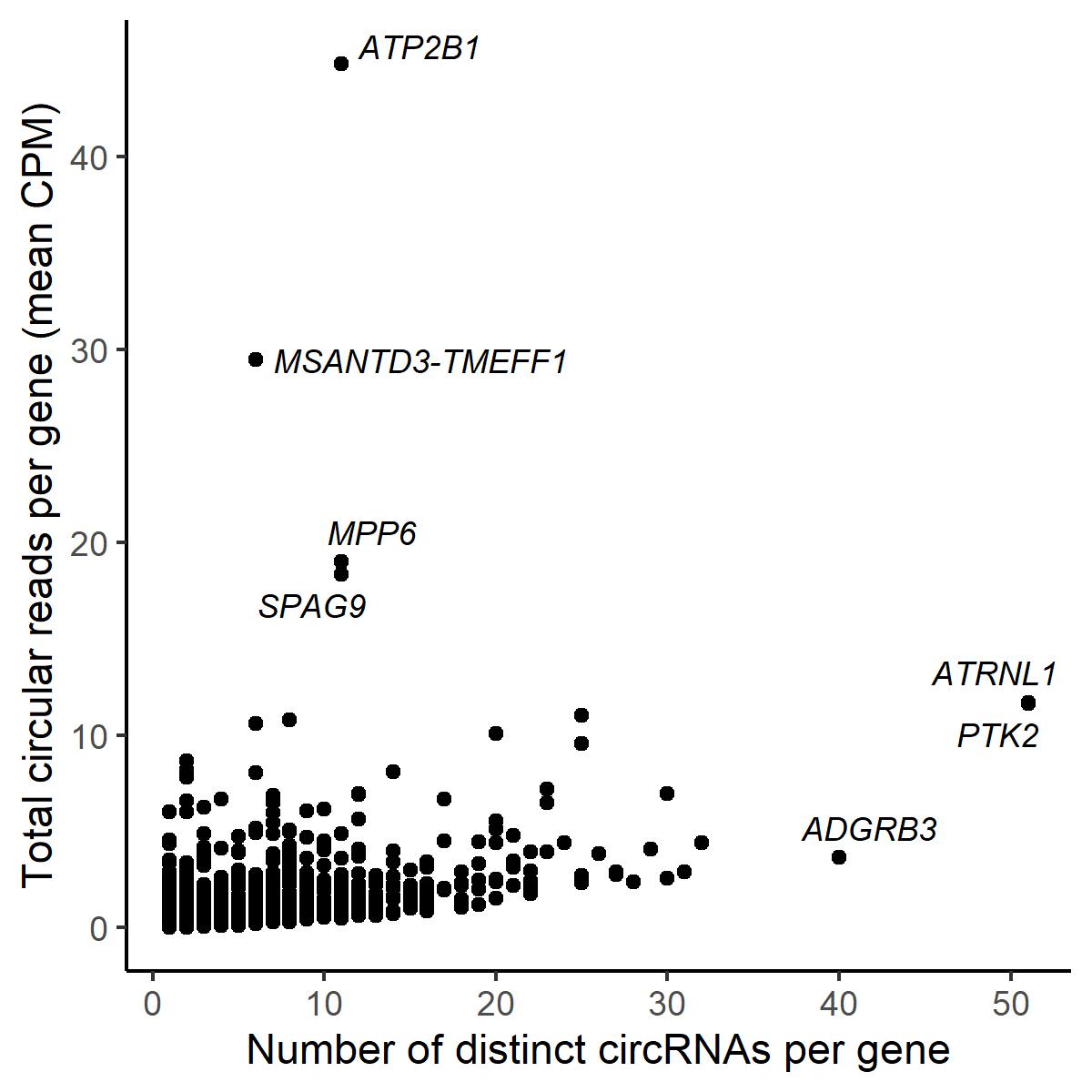

**Figure S1.** Comparison of the mean CPM of circular reads per gene with the number of distinct circRNAs per gene in the replication dataset.

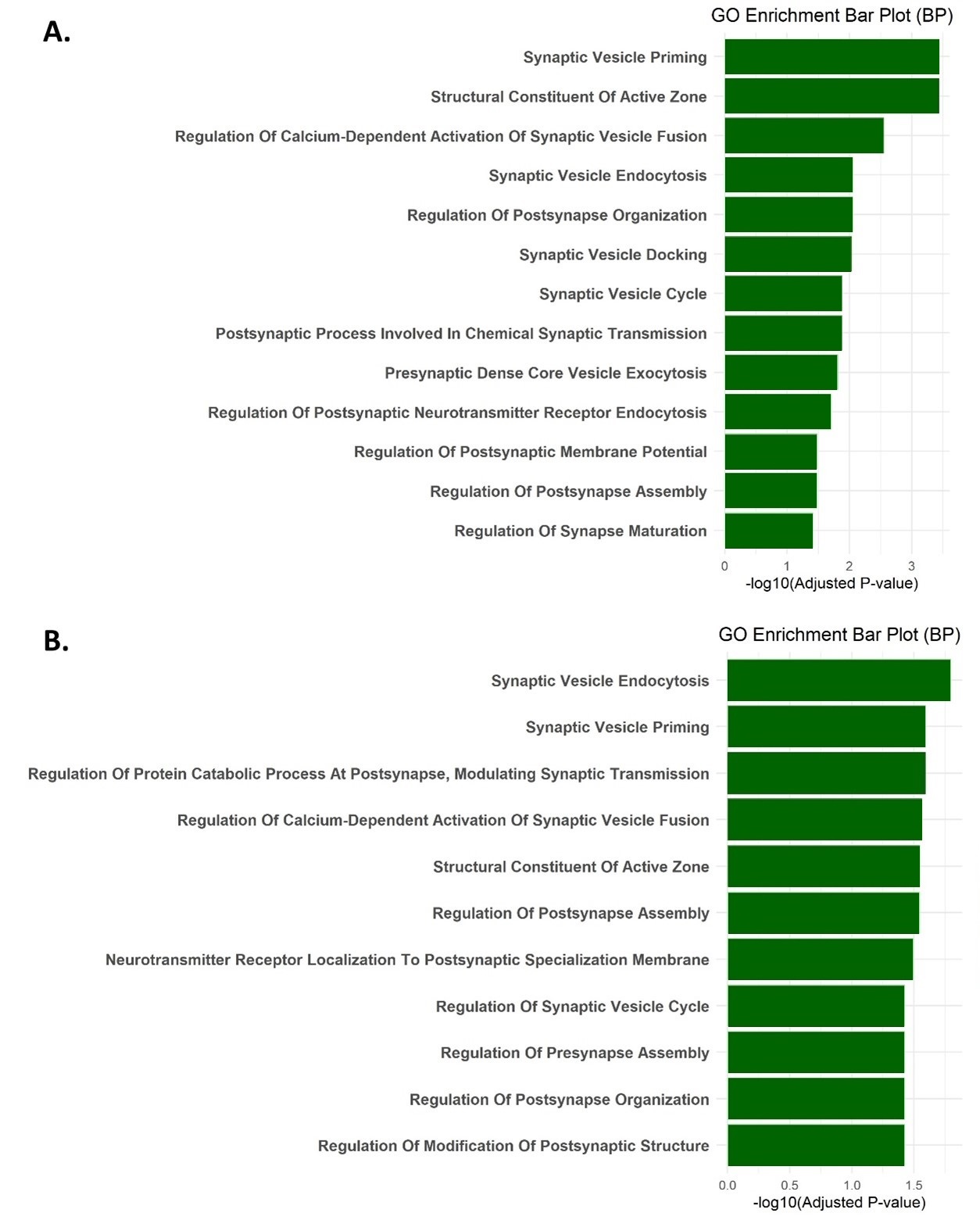

**Figure S2.** Functional enrichment analysis of the parental genes of circRNAs in modules M4 (A) and M5 (B). The bar plot shows enriched Gene Ontology (GO) terms related to synaptic processes (SynGO database), with the x-axis representing the significance level (-log10 of the adjusted p-value) and the y-axis showing the GO terms.

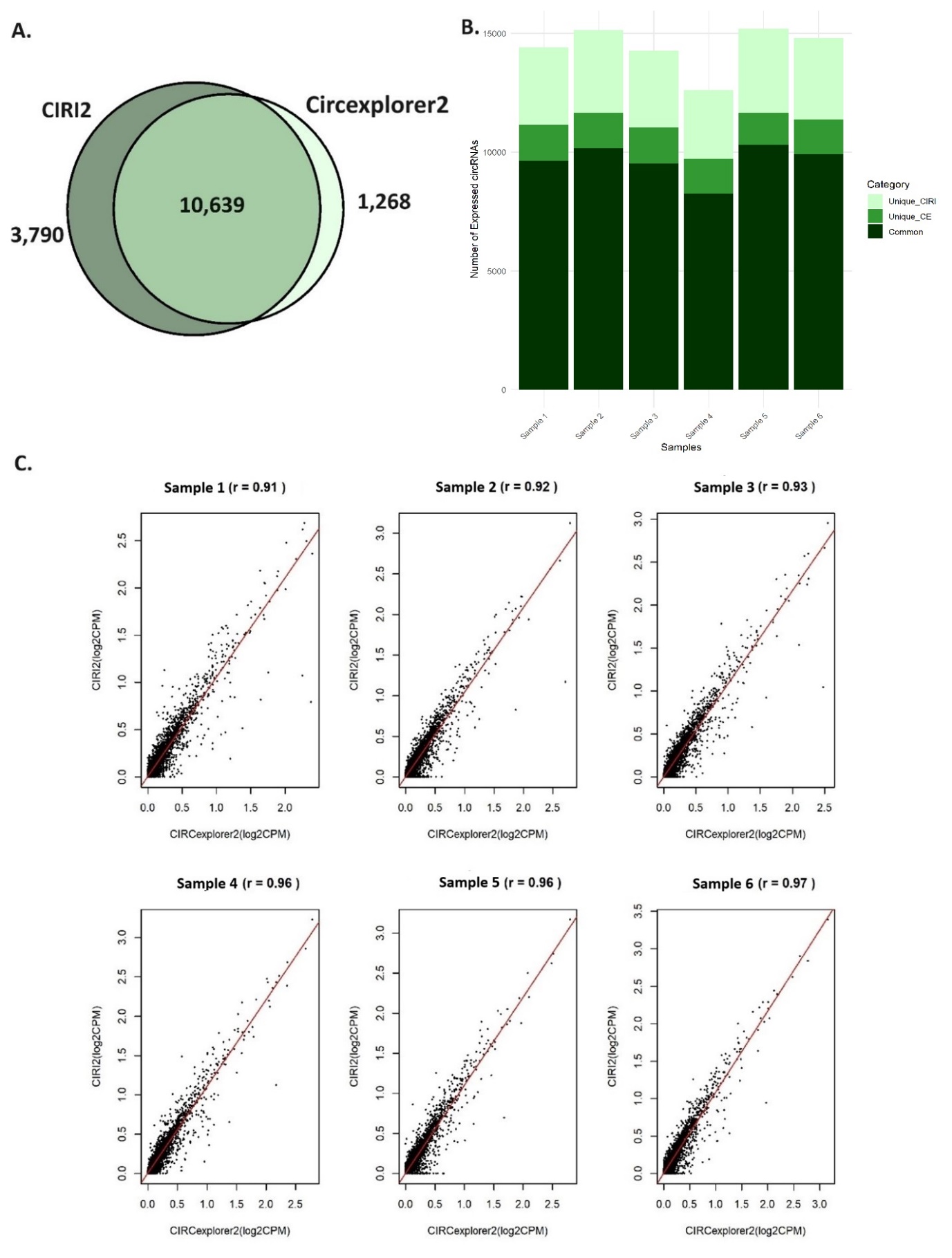

**Figure S3. A.** The Venn diagram illustrates the overlap in circRNA detection between CIRCexplorer2 and CIRI2 in discovery dataset. **B.** Number of circRNAs identified in 6 random samples by CIRCexplorer2 and CIRI2. **C.** The correlation of CPM values (log2 transformed) by CIRCexplorer2 and CIRI2 in 6 random samples.
